# CRYO-CEST: Non-invasive imaging of cryoprotectants using chemical exchange saturation transfer

**DOI:** 10.1101/2025.07.05.663058

**Authors:** Jan-Rüdiger Schüre, Moritz Zaiss, Arnd Dörfler, Alexander German

## Abstract

Chemical exchange saturation transfer (CEST) MRI at 7T was explored as a non-invasive method to detect and quantify common cryoprotectants used in organ cryopreservation. Phantom experiments demonstrated clear CEST signals from ethylene glycol, formamide, and dimethyl sulfoxide, with formamide showing the most sensitive concentration-dependent response. CEST imaging could thus provide a practical tool for monitoring cryoprotectant concentrations in organs.

## Introduction

Cryopreservation holds promise for transplantation medicine via improved utilization and immunological matching of organs [1]. Successful organ cryopreservation and transplantation has recently been achieved in rodents, using vitrification [2-5]. Vitrification enables ice-free cryopreservation by replacing high proportions of tissue water with polar solvents (cryoprotective agents, CPA) to avoid water crystallization at cryogenic temperatures [6]. CPA introduction into organs via perfusion of the vasculature has to be precisely controlled: The minimal CPA concentration needed to vitrify must be exceeded in all parts of the organs to preclude damage from ice-formation during cooling and rewarming. At the same time, CPA toxicity correlates with increasing CPA concentration, exposure temperature, and time [7]. Therefore, techniques for precisely determining the moment of sufficient, but not excessive cryoprotectant equilibration in all parts of an organ before cryopreservation for transplantation would be of great utility. Typically employed CPAs for vitrification include ethylene glycol (EG), dimethyl sulfoxide (DMSO), and formamide (FA) [8]. While non-invasive x-ray computed tomography imaging of DMSO is possible due to the radiodensity of its sulfur atom [9, 10], FA and EG are only composed of CHON elements.

## Methods

All three CPAs were analyzed individually. For this purpose, different amounts of concentrations (2, 4, 6, 8, 10, 12, 14, 16, 18, 20, 22 %w/v) were combined in 15ml phantom tubes with distilled water. Image acquisition was performed on a 7T system, by using a 3D snapshot CEST sequence with three different B1 levels (0.4, 0.6, 0.9 muT). CEST data were acquired from -4 to 4 ppm with 0.1 ppm sampling. To correct for B_0_ and B_1_ field inhomogeneities, WASABI [11] was employed to simultaneously obtain B_0_ and relative B_1_ maps. In order to calculate the MTRasym, an equidistant sampling was used for the Z-spectrum acquisition. An omega-plot [12] was calculated to determine the different concentrations in each tube retrospectively according to 1/MTR_asym_ vs. 1/ω_1_^2^. Since the slope of each line in the omega plot is inversely proportional to the concentration of the exchanging proton pool, the concentration can be directly inferred from the slope.

## Results

Figure 1. shows for FA and EG that the peaks in both the Z-spectrum and the MTRasym appear to correlate with the respective concentration. While FA has the advantage to be very sensitive with concentration changes, EG has the advantage to resonate near resonance frequency of water and thus have less contributors through the NOE, ranging from -1 to -5 ppm. However, in the case of DMSO, additional NOE appear to occur which interfere with an adequate evaluation by the MTRasym.

**Fig.1.**
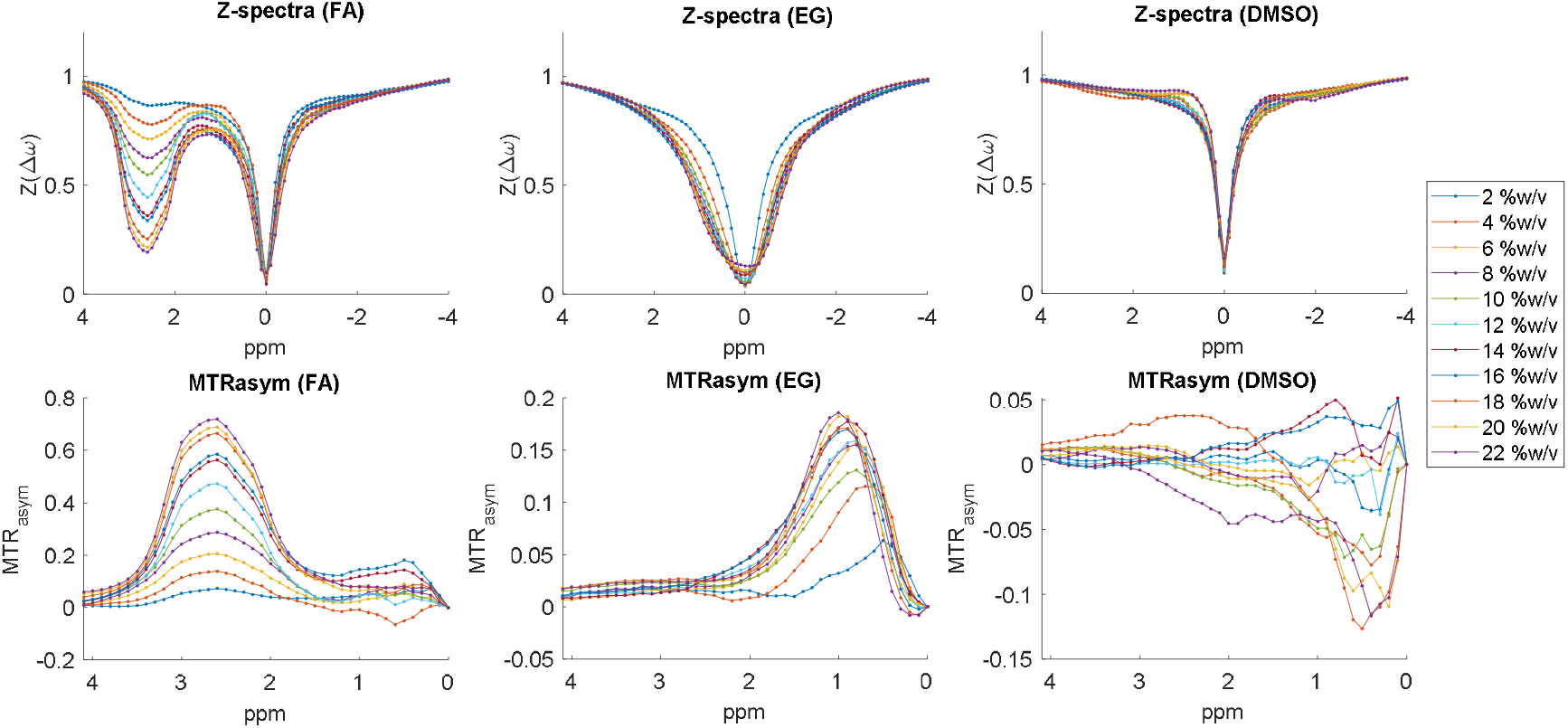
Calculation of the relative concentration based from the omega-plots in each individual phantom tube with 2, 4, 6, 8, 10, 12, 14, 16, 18, 20, 22 %w/v of each CPA. While Fa is most sensitive to concentration changes, EG occurs near the resonance frequency of water, making it more stable towards NOE effects in the MTRasym signal. DMSO shows a mixed contribution in MTRasym and seems less suitable.

Figure 2. shows the relative concentration using FA as an example, calculated from the slope of the omega plots at 2.6 ppm. Based on the known concentration of phantom tube 11 (22 % w/v), all other values were calculated relative to this reference concentration.

**Fig.2.**
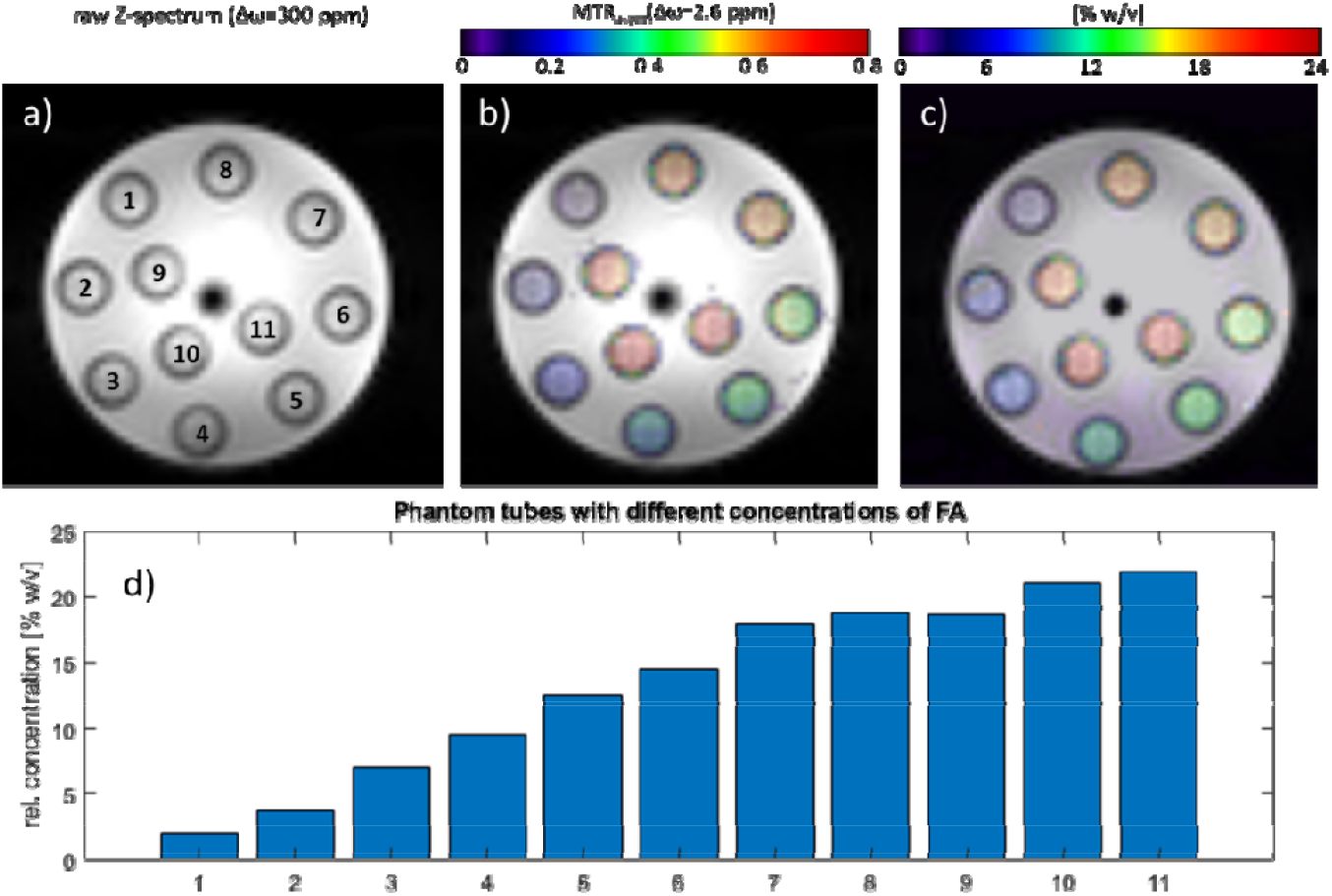
Illustration of a) the phantom and the tube reordering, b) the corresponding MTRasym at the resonance frequency of formamide at 2.6 ppm, c) the calculated FA concentration map and d) the corresponding bar plot, based on omega-plot data from the different B1 acquisitions. To obtain the relative concentrations, we calculated the values based on the phantom tube 11 with 22% w/v.

## Discussion

The in vitro experiment shows that CPAs can be detected by CEST imaging. Assuming that no further MT effects occur in the Z-spectrum, the relative concentration can be calculated from the MTRasym. It has been demonstrated that EG oscillates in close proximity to the resonance frequency of water. Consequently, it is less susceptible to interference from NOE factors. However, EG shows a slight drift of the amplitude in MTRasym. FA reacts even more sensitively to changes in concentration and could therefore be a promising CPA marker. Further studies on in vivo tissues such as porcine organs are planned to verify the effect by including further MT effects and performing CEST imaging before and after CPA loading.

## Conclusion

In conclusion, cryoprotectants can be detected using CEST imaging. This seems to be a promising approach to monitor CPA concentration during organ cryopreservation.

## References

1. Giwa, S., et al., The promise of organ and tissue preservation to transform medicine. Nature Biotechnology, 2017. 35(6): p. 530–542.

2. Chiu-Lam, A., et al., Perfusion, cryopreservation, and nanowarming of whole hearts using colloidally stable magnetic cryopreservation agent solutions. Sci Adv, 2021. 7(2).

3. Han, Z., et al., Vitrification and nanowarming enable long-term organ cryopreservation and life-sustaining kidney transplantation in a rat model. Nat Commun, 2023. 14(1): p. 3407.

4. Sharma, A., et al., Cryopreservation of Whole Rat Livers by Vitrification and Nanowarming. Ann Biomed Eng, 2023. 51(3): p. 566–577.

5. Wowk, B., et al., 27 MHz constant field dielectric warming of kidneys cryopreserved by vitrification. Cryobiology, 2024. 115: p. 104893.

6. Fahy, G.M., et al., Vitrification as an approach to cryopreservation. Cryobiology, 1984. 21(4): p. 407–26.

7. Fahy, G.M., Principles of Vitrification as a Method of Cryopreservation in Reproductive Biology and Medicine, in Fertility Preservation: Principles and Practice, J. Donnez and S.S. Kim, editors. 2021, Cambridge University Press: Cambridge. p. 49–66.

8. Fahy, G.M., et al., Improved vitrification solutions based on the predictability of vitrification solution toxicity. Cryobiology, 2004. 48(1): p. 22–35.

9. Bleisinger, N., et al., Me2SO perfusion time for whole-organ cryopreservation can be shortened: Results of micro-computed tomography monitoring during Me2SO perfusion of rat hearts. PLoS One, 2020. 15(9): p. e0238519.

10. Wang, S., et al., Viable Vitreous Grafts of Whole Porcine Menisci for Transplant in the Knee and Temporomandibular Joints. Advanced Healthcare Materials, 2024. 13(22): p. 2303706.

11. Schuenke P, et al., Simultaneous mapping of water shift and B_1_ (WASABI)-Application to field-Inhomogeneity correction of CEST MRI data. Magn Reson Med. 2017 Feb;77(2):571–580

12. Wu R, et al. Quantitative chemical exchange saturation transfer (qCEST) MRI - omega plot analysis of RF-spillover-corrected inverse CEST ratio asymmetry for simultaneous determination of labile proton ratio and exchange rate. NMR Biomed. 2015 Mar;28(3):376–83.

